# Far-red timing uncovers cultivar-dependent yield and bolting responses in vertical-farm spinach (Spinacia oleracea L.)

**DOI:** 10.64898/2026.07.10.737849

**Authors:** Christopher McGovern, Milena Adrio, Hadi Aliki, Ruth Vichos, Wayne Powell, Rajiv Sharma

**Affiliations:** Horticultural Innovation centre, School of Natural and Social Sciences, Scotland’s Rural College (SRUC), Kings Buildings, West Mains Road, Edinburgh EH9 3JG, UK

**Keywords:** vertical farming, controlled-environment agriculture, far-red light, shade avoidance, bolting, Spinacia oleracea, genotype × environment × management, light recipe

## Abstract

Far-red light (FR; 700-750 nm) is increasingly incorporated into controlled-environment lighting because it can improve photosynthetic efficiency when combined with comparatively shorter wavelengths. In long-day leafy crops such as spinach, however, FR may also promote the transition from vegetative to reproductive growth and thereby reduce marketable yield. Most studies have evaluated FR fraction, intensity or end-of-day exposure, whereas the developmental timing of FR has rarely been tested, particularly in spinach. Here, we evaluated six commercial spinach cultivars (Amador, Harp, Renegade, Responder, Rubino and Santa Cruz) in an indoor vertical farm under a common red-green-blue background (PPFD 260-264 µmol m⁻² s⁻¹, 12 h photoperiod, 24 °C) and four FR timing treatments: no FR (Control), FR throughout production (FullFR), FR during early development only (EarlyFR), and FR during late development only (LateFR). LateFR increased marketable fresh weight relative to Control (244 vs 224 g) and reduced flowering incidence, whereas far-red supplied during early development reduced fresh weight (158 g) and increased flowering. The magnitude of the timing response differed among cultivars: switching from EarlyFR to LateFR recovered 0 % fresh weight in Amador but 107 % in Renegade and Rubino, with the largest penalties occurring in otherwise bolt-resistant cultivars. EarlyFR also increased total chlorophyll and reduced the chlorophyll a:b ratio. These results show that FR response in spinach is strongly conditioned by developmental stage and cultivar. Although LateFR received more total far-red than EarlyFR, it behaved like the Control, indicating that the penalty was set by far-red timing rather than dose. Treatment differences in bolting and yield tracked an estimated phytochrome photostationary-state deficit during early development: a phytochrome-deficit model markedly outperformed a cumulative-dose model (ΔAIC = 441), and the deficit × cultivar interaction was strong (p < 0.001), with bolt-resistant cultivars losing most yield when far-red coincided with the early developmental window. We therefore propose that FR should be treated as a genotype-dependent management variable rather than as a fixed spectral input, with late application and bolt-resistant cultivars offering the most favourable combination for vertical-farm spinach production. Framed within the breeder’s equation, the close match between the trial and production environment and the scope for shorter breeding cycles indoors suggest that genotype and far-red timing can be optimised jointly to accelerate genetic gain.

## Introduction

In northern Europe and other high-latitude regions, fresh vegetable supply is increasingly affected by population growth, urbanisation, climate instability and disruption to long food, fertiliser and energy supply chains. These pressures have increased interest in local, season-independent production systems that can buffer seasonal shortages and reduce dependence on long-distance transport (Giradet, 2008; Sulaiman & Mirabi, 2023). Leafy vegetables are an important target because field production is highly seasonal and because warm, long-day conditions can accelerate flowering in cool-season crops.

Controlled-environment vertical farms (CEVFs) decouple crop production from season and latitude by replacing sunlight with light-emitting diodes (LEDs) and maintaining defined set-points for temperature, humidity, CO₂ and nutrition (Martínez-Moreno et al., 2024; Farhangi et al., 2025). These systems can increase land-use efficiency, shorten supply chains and reduce some chemical inputs such as Fertilizers application as well as pesticides (Despommier, 2011; Sulaiman & Mirabi, 2023). The principal constraint on indoor vertical farming remains the high operational cost, particularly electricity for artificial lighting. Rising energy prices in recent years, driven by the COVID-19 pandemic and subsequent disruptions to global energy markets, have further increased production costs (Despommier, 2011; Ferrarezi et al., 2024). Consequently, optimisation of indoor leafy-green production requires light recipes that improve marketable output per unit energy rather than simply maximise biomass.

Spinach (Spinacia oleracea L.) is well suited to vertical production in several respects: it is compact, fast growing, nutritionally dense and increasingly demanded in urban diets (Zou et al., 2020; Martínez-Moreno et al., 2024). It is also agronomically challenging because it is a long-day crop that bolts readily under photoperiods of approximately 13-15 h or under warm conditions. Bolting shifts growth from harvestable rosettes to reproductive stem elongation, reducing yield and quality traits (Chun et al., 2000; Semenova et al., 2023; Amini et al., 2025). Managing the balance between vegetative growth and reproductive transition is therefore central to indoor spinach production.

For plants, the red (R; 600-700 nm) to far-red (FR; 700-750 nm) ratio is a major signal of neighbouring vegetation. Chlorophyll absorption reduces R relative to FR within dense canopies, and plants interpret a low R:FR ratio as an indication of competition (Li et al., 2011; Wei, Wang & Yu, 2023). This signal is absorbed mainly through phytochrome B and PHYTOCHROME-INTERACTING FACTOR pathways and can induce shade-avoidance responses, including stem and petiole elongation, hyponasty, altered chlorophyll partitioning, and earlier flowering or bolting (Franklin & Whitelam, 2005; Casal, 2012; Paik & Huq, 2019). At the same time, FR photons can contribute efficiently to photosynthesis when combined with shorter wavelengths by balancing excitation between photosystems I and II (Zhen & Bugbee, 2020). Thus, FR may increase light-use efficiency but can also activate developmental responses that reduce spinach marketability and thus profitability.

Most work on FR in controlled environments has used model species or lettuce, and has applied FR continuously or as an end-of-day treatment (Li & Kubota, 2009; Kalaitzoglou et al., 2019; Tan et al., 2022). Spinach has been studied less extensively, and existing research has focused primarily on light quality (far-red fraction) and light quantity (photon flux density), while the influence of light duration (photoperiod) and the temporal distribution of far-red light within the photoperiod has received little attention (Lőrinc et al., 2019; Skabelund, Langenfeld & Bugbee, 2025). Skabelund et al. (2025) reported no morphological response of spinach to increasing far-red fraction across a wide range of photon fluxes, contrasting with responses reported in other leafy crops.

In this study, we tested FR timing as a management factor and evaluated its interaction with cultivar within a genotype × environment × management (G×E×M) framework. The objective was to determine how the timing of supplemental FR affects yield, leaf morphology, chlorophyll traits and bolting in spinach, and to identify whether cultivars differ in their sensitivity to FR timing. We hypothesised that FR would increase flowering probability and plant height, increase yield when applied at an appropriate stage, have a neutral or negative effect on chlorophyll content, and produce cultivar-dependent responses.

## Materials and methods

### Plant material and cultivation

Six commercial spinach cultivars (Amador, Harp, Renegade, Responder, Rubino and Santa Cruz; Moles Seeds Ltd, UK) were sown in 40-cavity propagation trays containing a 90:10 (v/v) coco-coir:vermiculite substrate. Trays were placed in an Intelligence Growth System (IGS) Growth Tower (model GT6) (https://www.intelligentgrowthsolutions.com/product/growth-towers) and germinated in darkness for seven days. Following uniform germination, trays were transferred to the light treatments. Two weeks after emergence, a nematode drench (Nemasys Biological Fruit and Veg Protection) and sticky traps were applied for sciarid fly control. Nutrient solution (90 L; composition in Table 2) was supplied by ebb-and-flow every two to three days at pH 5.8 and electrical conductivity 2.0 mS cm⁻¹. Air temperature, relative humidity, CO₂ concentration and photoperiod were maintained at 24 °C, 60 %, 800 ppm and 12 h, respectively.

### Light treatments and timing

All treatments received the same red-green-blue background spectrum at a mean PPFD of 260-264 µmol m⁻² s⁻¹ and an approximate daily light integral of 11.3 mol m⁻² d⁻¹ over 12 h. FR was supplied additively at 26 µmol m⁻² s⁻¹, equivalent to 10 % of PPFD. Four timing treatments were imposed: Control (no FR), FullFR (FR throughout the 32-day cycle), EarlyFR (FR during days 0-12 only) and LateFR (FR from the early/mid transition to harvest). Because FR was added to a constant PAR background, total photon flux differed between FR and non-FR phases while PAR remained constant. FR is therefore interpreted as a supplemental signal rather than a replacement for PAR. Treatment recipes are summarised in Table 1.

**Table 1.** Far-red timing treatments. All treatments received the same red-green-blue background (PPFD 260-264 µmol m⁻² s⁻¹; 12 h photoperiod; DLI approximately 11.1-11.4 mol m⁻² d⁻¹). Far-red, where applied, was supplied additively at 26 µmol m⁻² s⁻¹ (10 % of PPFD).

| Treatment | Far-red window | Days with FR | FR (μmol m<sup>-2</sup> s<sup>-1</sup>) | Total cycle (d) |
| --- | --- | --- | --- | --- |
| Control | None — RGB background only | — | 0 | 32 |
| FullFR | Whole cycle | 0–32 | 26 | 32 |
| EarlyFR | Early development only | 0–12 | 26 | 32 |
| LateFR | Late development only | 13–31 | 26 | 31 |

**Table 2.** Nutrient solution composition (per 10 L of stock).

| Stock | Component | g per 10 L |
| --- | --- | --- |
| A | Calcium nitrate | 960 |
| A | Potassium nitrate | 28 |
| A | Iron EDDHA 6% | 75 |
| B | Potassium nitrate | 462 |
| B | Magnesium sulphate | 350 |
| B | Monopotassium phosphate | 160 |
| B | Magnesium nitrate | 60 |
| B | Manganese sulphate | 3.6 |
| B | Disodium octaborate (21% B) | 2.4 |
| B | Zinc sulphate | 1.5 |
| B | Copper sulphate | 0.4 |
| B | Sodium molybdate | 0.2 |

### Experimental design

Each treatment consisted of 30 propagation trays, with five replicate trays per cultivar (n = 5 trays per cultivar per treatment; n = 30 trays per treatment). Individual plants within trays (target 40 plants per tray) were treated as sub-samples.

### Data collection

Plant height, leaf length (LL) and leaf width (LW) were measured on five randomly selected plants per tray at 17 days (T1) and 32 days (T2) after emergence. Height was measured from the substrate surface to the newest apical growth; at T2, where bolting occurred, height was measured to the topmost growth or flower. This measure therefore partly includes bolting and is interpreted accordingly in the Discussion. Aerial fresh weight was recorded immediately after harvest and samples were then oven-dried at 80 °C for 72 h to determine dry weight. Chlorophyll was extracted from random two leaf samples per tray by boiling in at least 99 % ethanol at 85 °C for 15 min, followed by absorbance readings at 647 and 664 nm. Total chlorophyll and chlorophyll a:b ratio were calculated per sample and are reported as relative values rather than per unit leaf mass. Flowering incidence was recorded as the number of flowering or bolting plants divided by the total number of plants per tray at harvest.

### Statistical analysis

Analyses were conducted in R version 4.3.3 (https://www.r-project.org/). Because light treatments were applied at tray level, tray was the experimental unit for treatment effects. Continuous traits were analysed using linear mixed-effects models (using lme4::lmer function) with treatment and cultivar as fixed effects. Weight and chlorophyll traits were analysed at tray level, whereas morphology traits were analysed at plant level with tray included as a random effect to account for sub-sampling. Sum-to-zero contrasts were used so that Type III Wald F tests (car::Anova with Kenward-Roger denominator degrees of freedom) tested main effects in the presence of interactions. Height was analysed across both sampling times using a Time × treatment × cultivar model. Significant effects were followed by estimated marginal means (emmeans) and Tukey-adjusted pairwise comparisons, or Sidak-adjusted comparisons for the binomial model. Flowering incidence was analysed using a binomial generalised linear mixed model (lme4::glmer), and the interaction was tested by likelihood-ratio test. Fresh weight was also analysed with flowering proportion included as a covariate. Significance was assessed at p < 0.05. Figures, compact-letter displays and the consolidated model summary (Table 3) were generated from a single reproducible R script applied to the master dataset. To indicate the relative contribution of each model term, the explained variation was additionally partitioned among the treatment (T), cultivar (V) and T × V terms using the exact Type III sums of squares from the split-plot decomposition (cultivar as the whole-plot factor and far-red timing as the sub-plot factor), each expressed as a percentage of the summed T, V and T × V components (Table 4); flowering incidence was partitioned analogously from the analysis of deviance of the binomial model. A quantitative-genetic summary of the phenotypic variation observed across cultivars, including broad-sense heritability estimates and projected genetic gains under different selection scenarios, is provided in Supplementary Text S1 and Table S6.

**Table 3.** Consolidated summary of fixed-effect tests. Values are Type III Wald F statistics with Kenward-Roger denominator degrees of freedom, except for flowering, where the row reports the GLMM likelihood-ratio χ². T = far-red timing treatment; V = cultivar.

| Response | Effect | F / $\chi^2$ | df | df (resid.) | p |
| --- | --- | --- | --- | --- | --- |
| Fresh weight (g) | T | 9.42 | 3 | 72 | <0.001 |
| Fresh weight (g) | V | 8.56 | 5 | 24 | <0.001 |
| Fresh weight (g) | T × V | 1.37 | 15 | 72 | 0.184 |
| Dry weight (g) | T | 5.06 | 3 | 72 | 0.003 |
| Dry weight (g) | V | 13.70 | 5 | 24 | <0.001 |
| Dry weight (g) | T × V | 1.86 | 15 | 72 | 0.043 |
| Leaf length T2 (cm) | T | 49.47 | 3 | 552 | <0.001 |
| Leaf length T2 (cm) | V | 16.54 | 5 | 24 | <0.001 |
| Leaf length T2 (cm) | T × V | 3.68 | 15 | 552 | <0.001 |
| Leaf width T2 (cm) | T | 29.24 | 3 | 552 | <0.001 |
| Leaf width T2 (cm) | V | 25.71 | 5 | 24 | <0.001 |
| Leaf width T2 (cm) | T × V | 2.91 | 15 | 552 | <0.001 |
| Leaf length:width T2 | T | 16.58 | 3 | 552 | <0.001 |
| Leaf length:width T2 | V | 100.99 | 5 | 24 | <0.001 |
| Leaf length:width T2 | T × V | 2.40 | 15 | 552 | 0.002 |
| Total chlorophyll (ug/mL) | T | 17.29 | 3 | 72 | <0.001 |
| Total chlorophyll (ug/mL) | V | 0.62 | 5 | 24 | 0.684 |
| Total chlorophyll (ug/mL) | T × V | 1.13 | 15 | 72 | 0.348 |
| Chlorophyll a:b ratio | T | 16.85 | 3 | 72 | <0.001 |
| Chlorophyll a:b ratio | V | 0.75 | 5 | 24 | 0.591 |
| Chlorophyll a:b ratio | T × V | 0.71 | 15 | 72 | 0.769 |
| Plant height (cm) | Time | 3044.53 | 1 | 1128 | <0.001 |
| Plant height (cm) | T | 46.45 | 3 | 1128 | <0.001 |
| Plant height (cm) | V | 68.82 | 5 | 24 | <0.001 |
| Plant height (cm) | Time × T | 13.02 | 3 | 1128 | <0.001 |
| Plant height (cm) | Time × V | 279.60 | 5 | 1128 | <0.001 |
| Plant height (cm) | T × V | 3.49 | 15 | 1128 | <0.001 |
| Plant height (cm) | Time × T × V | 2.38 | 15 | 1128 | 0.002 |
| Flowering-adj. fresh weight (g) | Flowering (cov.) | 5.60 | 1 | 93.6 | 0.020 |
| Flowering-adj. fresh weight (g) | T | 11.80 | 3 | 75.2 | <0.001 |
| Flowering-adj. fresh weight (g) | V | 1.73 | 5 | 28 | 0.160 |
| Flowering-adj. fresh weight (g) | T × V | 1.43 | 15 | 71.6 | 0.158 |
| Flowering (GLMM, Chisq) | T × V | 173.51 | 15 | — | <0.001 |

**Table 4.** Partition of explained variation among fixed-effect terms. Values are the percentage of the summed treatment (T), cultivar (V) and T × V variation attributable to each term, computed from the exact Type III sums of squares of the split-plot models underlying Table 3 (cultivar as the whole-plot factor, far-red timing as the sub-plot factor). For plant height, the time main effect and its interactions are excluded so that the three terms are comparable across traits. ‡ Flowering incidence is partitioned from the analysis of deviance of the binomial model (percentage of explained deviance; the split between the two main effects is averaged over entry order).

| Response | Treatment, T (%) | Cultivar, V (%) | T × V (%) |
| --- | --- | --- | --- |
| Fresh weight (g) | 30 | 48 | 22 |
| Dry weight (g) | 13 | 64 | 23 |
| Leaf length, T2 (cm) | 31 | 57 | 12 |
| Leaf width, T2 (cm) | 12 | 82 | 6 |
| Leaf length:width, T2 | 6 | 89 | 5 |
| Total chlorophyll | 71 | 6 | 23 |
| Chlorophyll a:b ratio | 77 | 7 | 16 |
| Plant height (cm)† | 6 | 92 | 2 |
| Flowering incidence‡ | 15 | 79 | 6 |

To test whether the timing effect reflected phytochrome state rather than far-red dose, we estimated the phytochrome photostationary state (PSS, φ = Pfr/Ptot) for each treatment from the programmed channel photon fluxes (red ∼660 nm, far-red ∼731 nm, with minor blue and green contributions), following φ = Σ Nλ σrλ / Σ Nλ (σrλ + σfrλ) (Sager et al., 1988). Because independent spectroradiometric scans were not recorded, φ is an estimate; the channel cross-sections were calibrated to reproduce Sager-based PSS values reported for fully specified recipes by Lanoue et al. (2022), which they were recovered to within 0.001 RMS. For each treatment we integrated the Pfr-deficit (φref − φ, where φref is the far-red-free reference state) over an early-leaf competence window (days 0–12), motivated by evidence that spinach registers photoperiodic floral signals during early transplant development (Chun et al., 2000). Plant-level bolting (binomial) and fresh weight were then related to the early-window Pfr-deficit and, separately, to cumulative far-red dose, using mixed models with the split-plot random structure; the non-nested deficit and dose models were compared by AIC. A 2000-draw cross-section sensitivity analysis confirmed that the treatment grouping was invariant to coefficient choice (100% of draws). Because the early-window deficit took two distinct values, cultivar reaction norms are two-level contrasts.

## Results

### Flowering incidence

Flowering incidence showed a strong treatment × cultivar interaction (likelihood-ratio test, χ²(15) = 173.5, p < 0.001; Fig. 1). Across cultivars, EarlyFR (73 %) and FullFR (71 %) increased flowering relative to Control (50 %) and LateFR (47 %). Cultivars separated into two broad groups. Amador, Harp and Rubino flowered heavily in all treatments, including Control (69-96 %), whereas Renegade, Responder and Santa Cruz showed low flowering under Control and LateFR (8-24 %) but increased sharply under EarlyFR and FullFR. Some cultivar-treatment combinations reached 100 % flowering, including Harp under LateFR. LateFR produced the lowest flowering incidence in five of six cultivars. These results support the hypothesis that FR can increase flowering, but show that the effect depends strongly on timing and cultivar.

**Figure 1.**
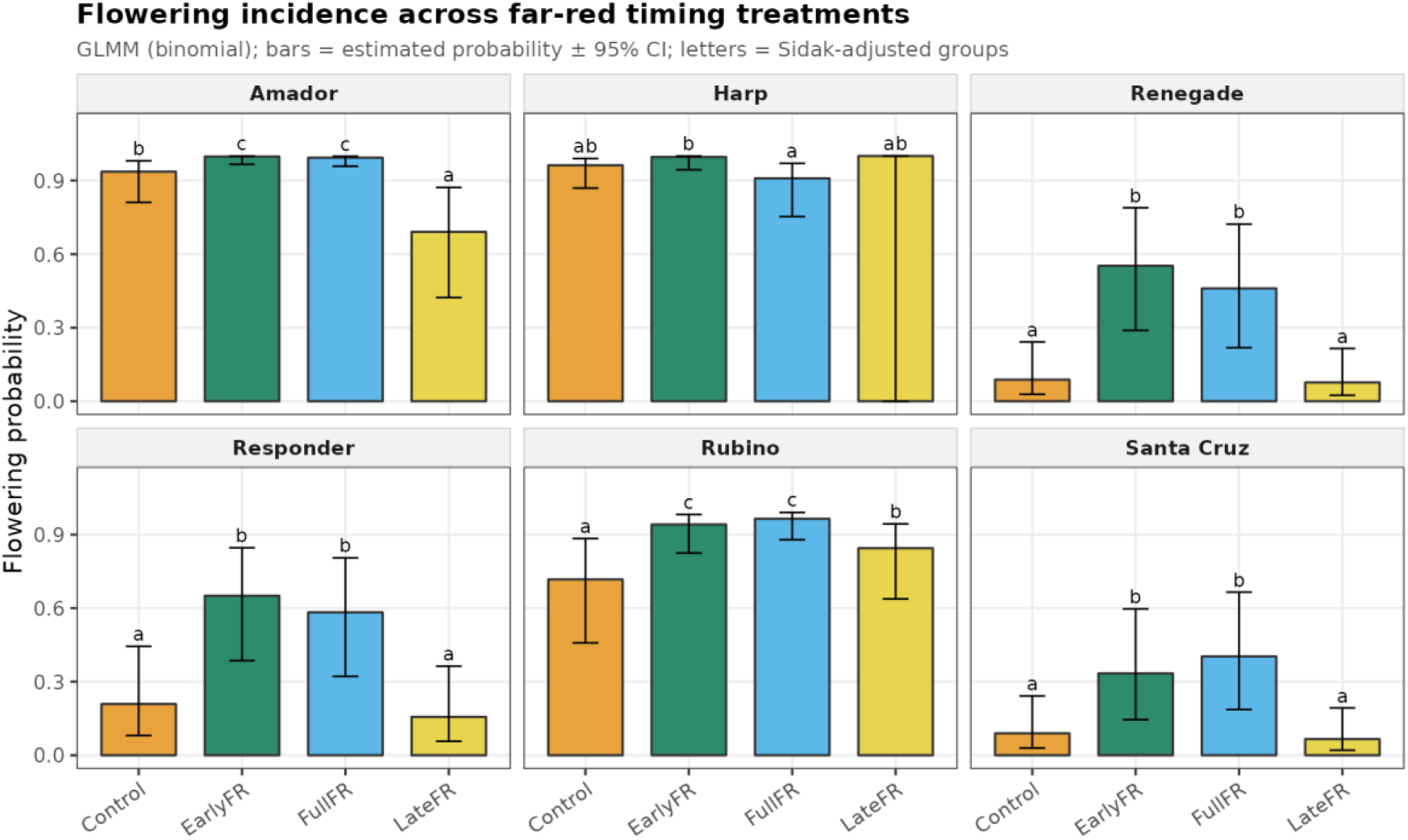
Flowering incidence (%) at harvest across far-red timing treatments, faceted by cultivar. Bars represent GLMM-estimated flowering probabilities ± 95 % CI; letters indicate Sidak-adjusted significance groups. Early and continuous far-red increased flowering, particularly in cultivars that remained comparatively bolt-resistant under Control and LateFR.

### Marketable yield and dry biomass

Fresh weight differed significantly among treatments (F(3,72) = 9.42, p < 0.001) and cultivars (F(5,24) = 8.56, p < 0.001), whereas the treatment × cultivar interaction was weak and not significant (F(15,72) = 1.37, p = 0.18). Treatment marginal means separated under Tukey comparison, with LateFR producing the greatest fresh weight (244 g, group c) and EarlyFR the lowest (158 g, group a); Control (224 g, group bc) and FullFR (192 g, group ab) were intermediate (Fig. 2a). Dry weight showed a significant treatment × cultivar interaction (F(15,72) = 1.86, p = 0.043), with cultivar explaining more variation than treatment (Fig. 2b). When flowering proportion was included as a covariate, both flowering (F(1,93.6) = 5.60, p = 0.020) and treatment (F(3,75.2) = 11.80, p < 0.001) remained significant. The yield advantage of LateFR over EarlyFR and FullFR was therefore not explained solely by differences in flowering. These results support the hypothesis that FR can increase yield when supplied late in development.

**Figure 2.**
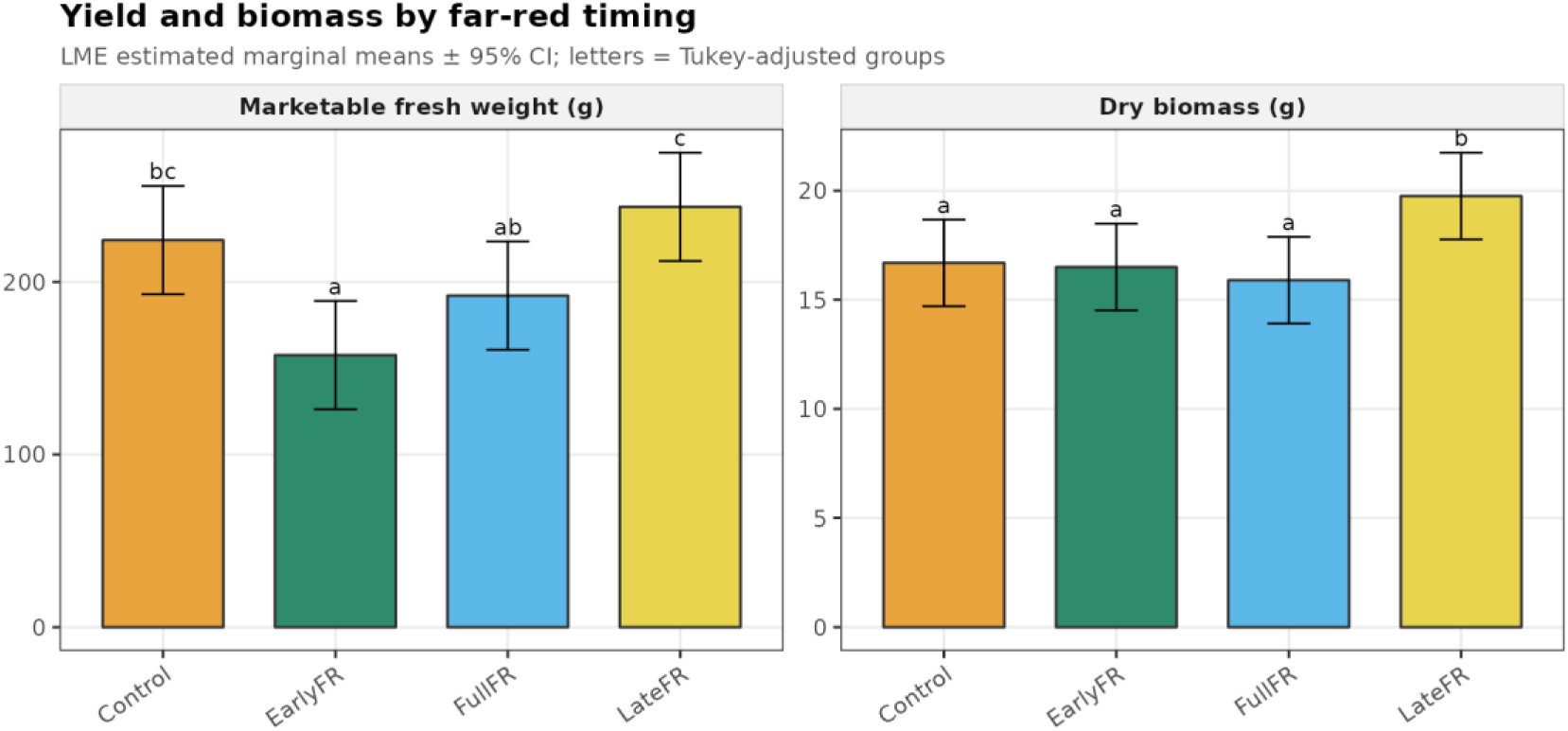
(a) Marketable fresh weight and (b) dry biomass by far-red timing treatment. Values are LME estimated marginal means ± 95 % CI; letters indicate Tukey groups. Late far-red produced the highest fresh weight, whereas early far-red produced the lowest.

### The genotype × timing interaction (G×E×M)

The yield response to FR timing was strongly cultivar-dependent (Fig. 3). Expressed as the fresh-weight gain from switching FR from EarlyFR to LateFR, recovery ranged from 0 % in Amador to 37 % in Harp, 67 % in Responder, 91 % in Santa Cruz and 106-107 % in Renegade and Rubino. When cultivars were grouped by bolting behaviour, the timing response was most pronounced in the bolt-resistant group. These cultivars maintained low flowering under Control and LateFR but lost substantial fresh weight under EarlyFR, consistent with early FR inducing bolting in genotypes that otherwise avoided it. By contrast, the bolt-prone group flowered heavily across treatments and showed a smaller relative timing response. These data indicate that FR timing acts as a management variable whose agronomic value depends on genotype, supporting the G×E×M hypothesis.

**Figure 3.**
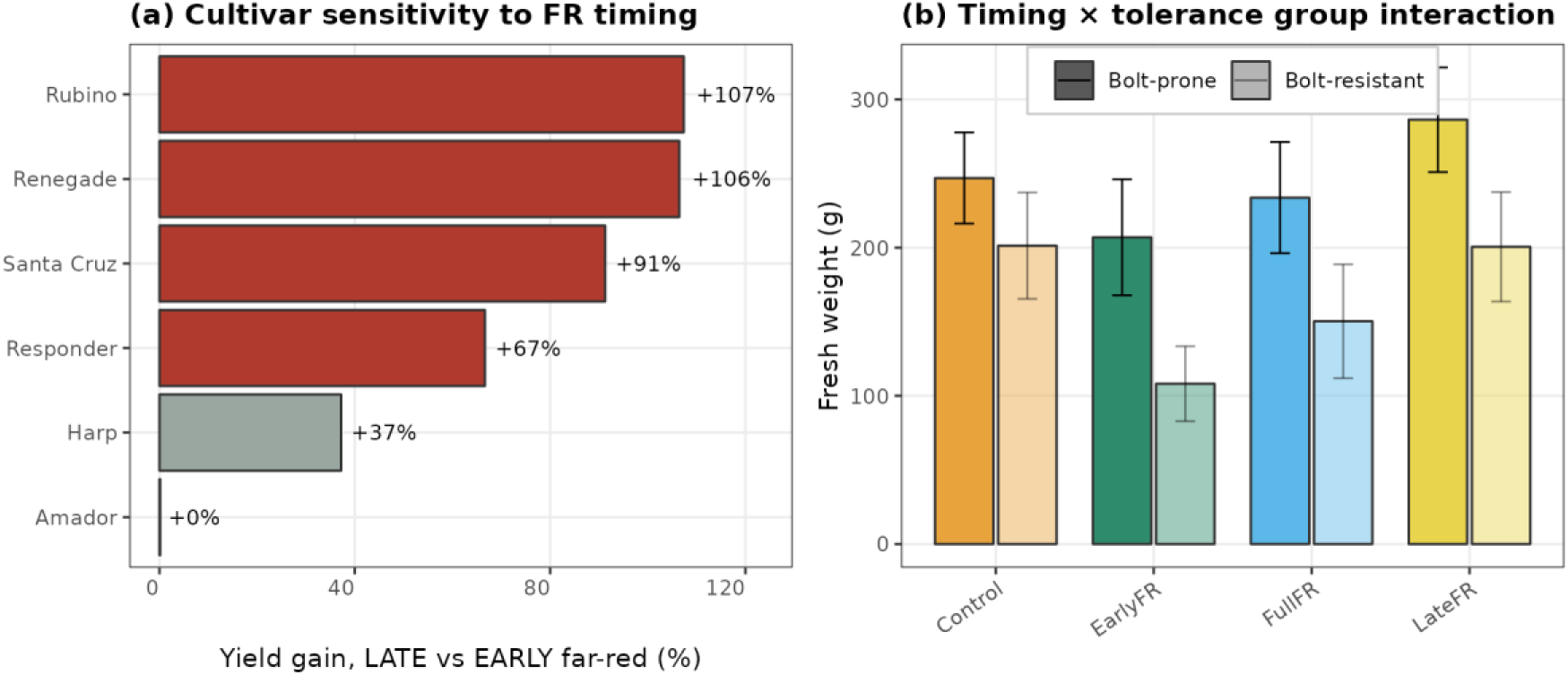
Genotype × environment × management interaction. (a) Fresh-weight gain (%) from switching far-red from early to late application, by cultivar; highlighted bars indicate timing gains greater than 50 %. (b) Mean fresh weight by treatment for bolt-prone and bolt-resistant cultivar groups. Bolt-resistant cultivars showed the largest penalty under EarlyFR and recovered under LateFR.

### Far-red timing acts through early-window phytochrome state, not dose

The treatments dissociated far-red timing from far-red dose. Integrating the estimated Pfr-deficit over the early-leaf window rendered EarlyFR and FullFR equivalent (≈ 0.37 φ·d) and LateFR equivalent to the Control (≈ 0 φ·d), despite a near three-fold range in cumulative far-red dose (EarlyFR 13.5, LateFR 21.3, FullFR 35.9 mol m⁻²; Fig. 4A,B). Bolting followed early-window phytochrome state rather than dose: a binomial mixed model in which bolting depended on the early-window Pfr-deficit markedly outperformed an otherwise identical model based on cumulative far-red dose (ΔAIC = 441 in favour of the deficit model), and LateFR — which delivered more total far-red than EarlyFR — patterned with the Control rather than with the far-red treatments (Fig. 4C). Marketable yield showed the same signature: across treatment means, fresh weight was strongly and negatively associated with the early-window deficit (r = −0.90) but essentially unrelated to far-red dose (r = −0.14), with EarlyFR yielding least despite receiving the least far-red (Fig. 4D).

**Figure 4.**
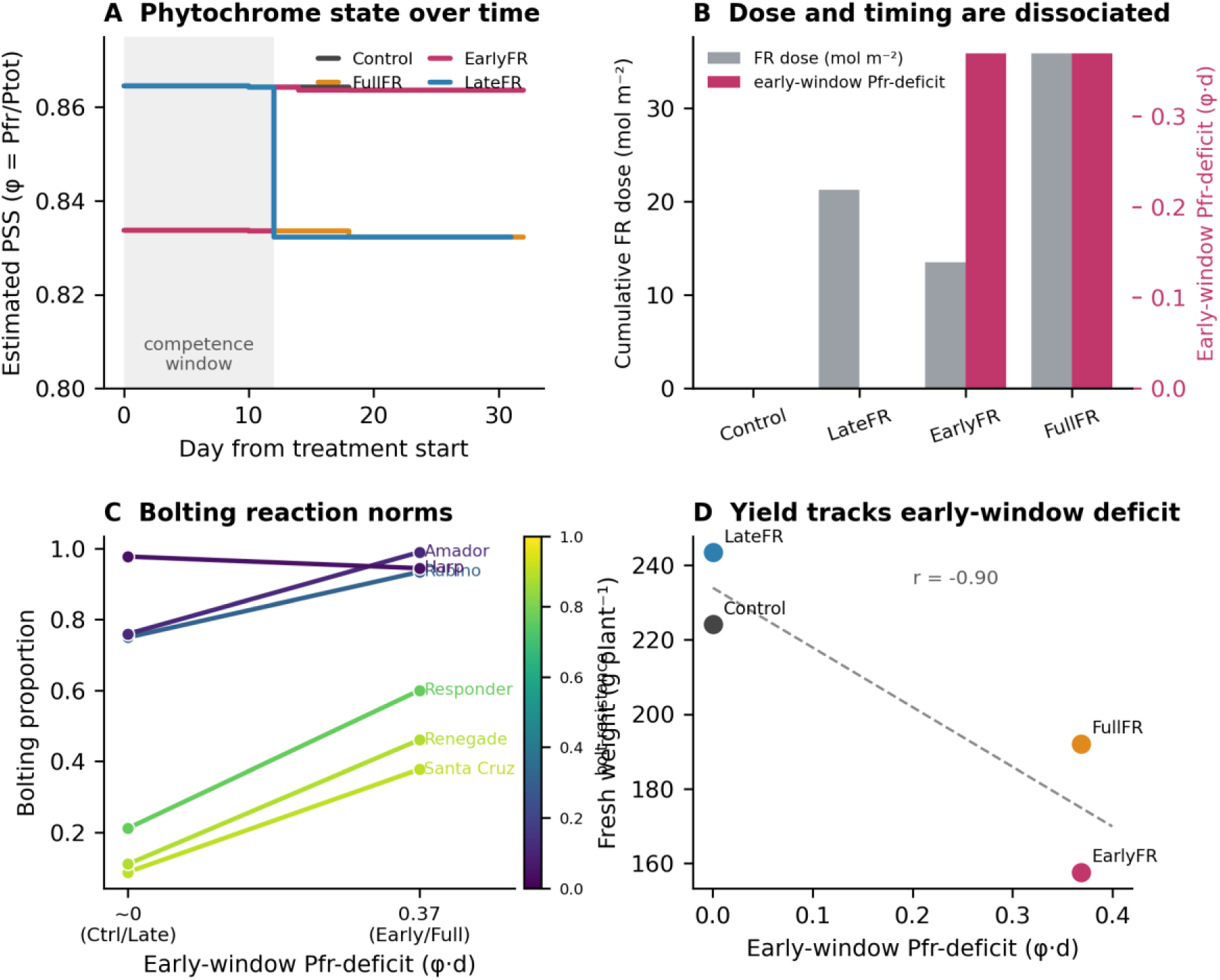
Far-red timing acts through early-window phytochrome state, not dose. (A) Estimated phytochrome photostationary state (φ) over the production cycle for each treatment; shading marks the assumed early-leaf competence window (days 0–12). (B) Cumulative far-red dose and integrated early-window Pfr-deficit per treatment, showing that EarlyFR carries the least dose yet the same early-window deficit as FullFR. (C) Cultivar reaction norms for bolting against the early-window Pfr-deficit (a two-level contrast), coloured by bolting tolerance. (D) Treatment-mean fresh weight against early-window Pfr-deficit (r = −0.90). φ values are estimates calibrated to published Sager-based PSS (Lanoue et al., 2022).

The bolting response to the early-window deficit was strongly cultivar-dependent (deficit × cultivar interaction, Wald χ² = 69.4, df = 5, p = 1.4 × 10⁻¹³; deficit main effect χ² = 50.3, p = 1.3 × 10⁻¹²). Under Control and LateFR the cultivars spanned the full range from near-obligate bolting (Harp, Amador) to strong tolerance (Santa Cruz, Renegade; Control bolting ≈ 0.10–0.12). Early-window far-red largely abolished this contrast: in the bolt-tolerant cultivars, bolting rose from a mean of 0.14 without early-window far-red to 0.48 with it, whereas the bolt-prone cultivars were already near ceiling and changed little (Fig. 4C). The effect is therefore best described as an erosion of the bolting tolerance that distinguishes resistant genotypes, rather than as a uniform sensitivity gradient; because bolt-prone cultivars approach the response ceiling, the per-cultivar sensitivity does not scale monotonically with tolerance on the latent scale (Spearman ρ = 0.37, p = 0.50, n = 6).

### Plant height

Height was analysed across T1 (day 17) and T2 (harvest) using a Time × treatment × cultivar model. Time explained the largest component of variation, as expected for a growth trait (F(1,1128) = 3044.5, p < 0.001), and all tested terms were significant: treatment (F(3,1128) = 46.5, p < 0.001), cultivar (F(5,24) = 68.8, p < 0.001), Time × cultivar (F(5,1128) = 279.6, p < 0.001), Time × treatment (F(3,1128) = 13.0, p < 0.001), treatment × cultivar (F(15,1128) = 3.49, p < 0.001) and the three-way interaction (F(15,1128) = 2.38, p = 0.002). At T1, treatments did not differ within cultivars (Fig. 5, upper row). By T2, FullFR and LateFR produced the tallest plants, averaging approximately 35 % and 32 % above Control, respectively, and the bolt-prone cultivars were tallest overall. These results support the hypothesis that FR increases height, although T2 height partly reflects bolting in susceptible cultivars.

**Figure 5.**
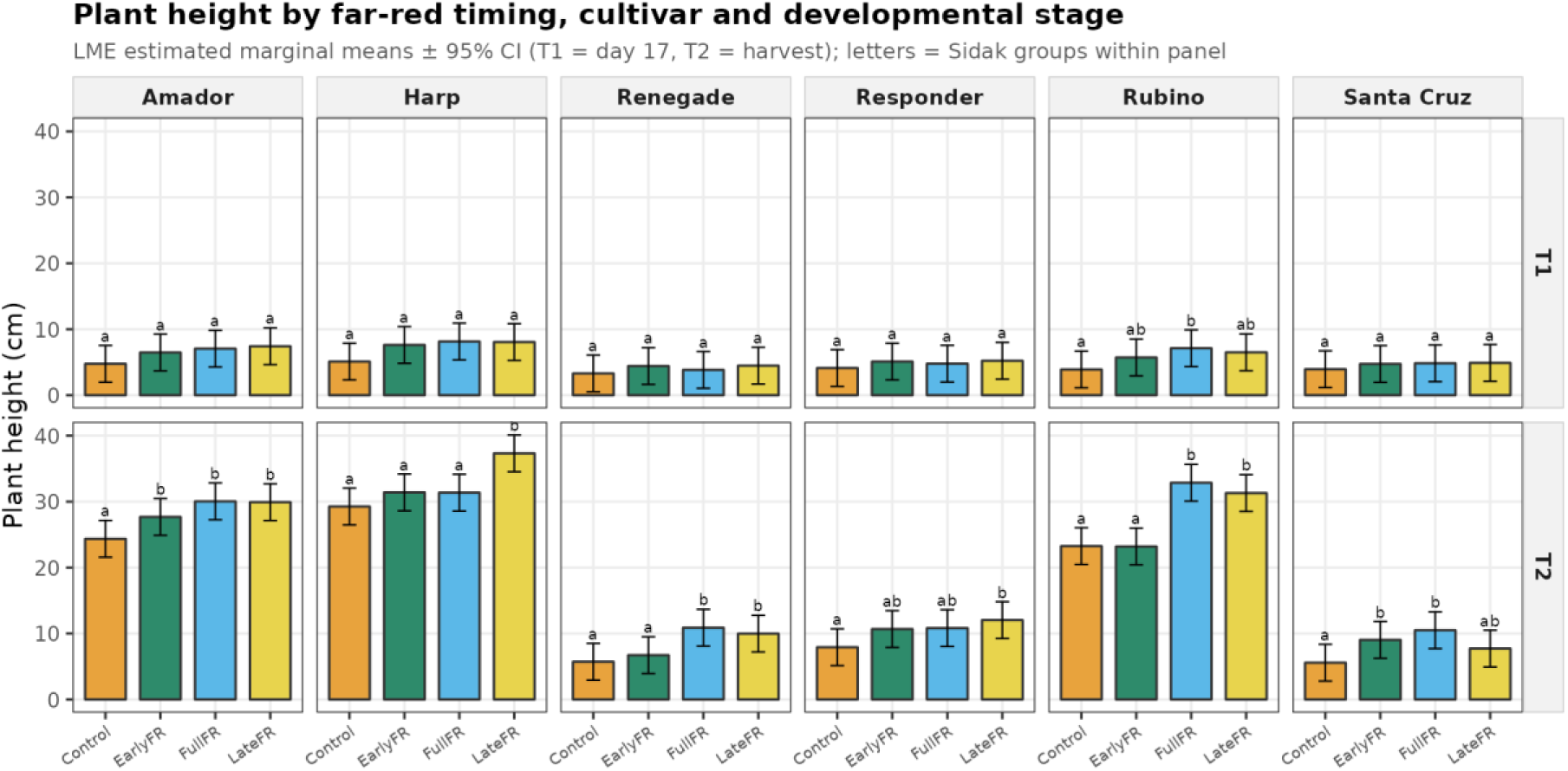
Plant height by far-red timing, cultivar and developmental stage (T1 = day 17, upper row; T2 = harvest, lower row). Bars represent LME estimated marginal means ± 95 % CI; letters indicate Sidak-adjusted groups within each panel. Treatment differences were negligible at T1 but evident by T2.

### Leaf morphology

Leaf length differed among treatments (F(3,552) = 49.5, p < 0.001), with Control and LateFR producing longer leaves than EarlyFR and FullFR. Cultivar (F(5,24) = 16.5, p < 0.001) and the treatment × cultivar interaction (F(15,552) = 3.68, p < 0.001) were also significant. Leaf width showed a similar response (treatment F(3,552) = 29.2, p < 0.001; cultivar F(5,24) = 25.7, p < 0.001; interaction F(15,552) = 2.91, p < 0.001), with the widest leaves under Control. The leaf length-to-width ratio increased under FR and was highest under LateFR (treatment F(3,552) = 16.6, p < 0.001; cultivar F(5,24) = 101.0, p < 0.001; interaction F(15,552) = 2.40, p = 0.002; Fig. 6). This pattern is consistent with shade-avoidance elongation, particularly under late FR. Because leaf and petiole were measured together, the ratio reflects combined leaf-and-petiole elongation rather than lamina shape alone.

**Figure 6.**
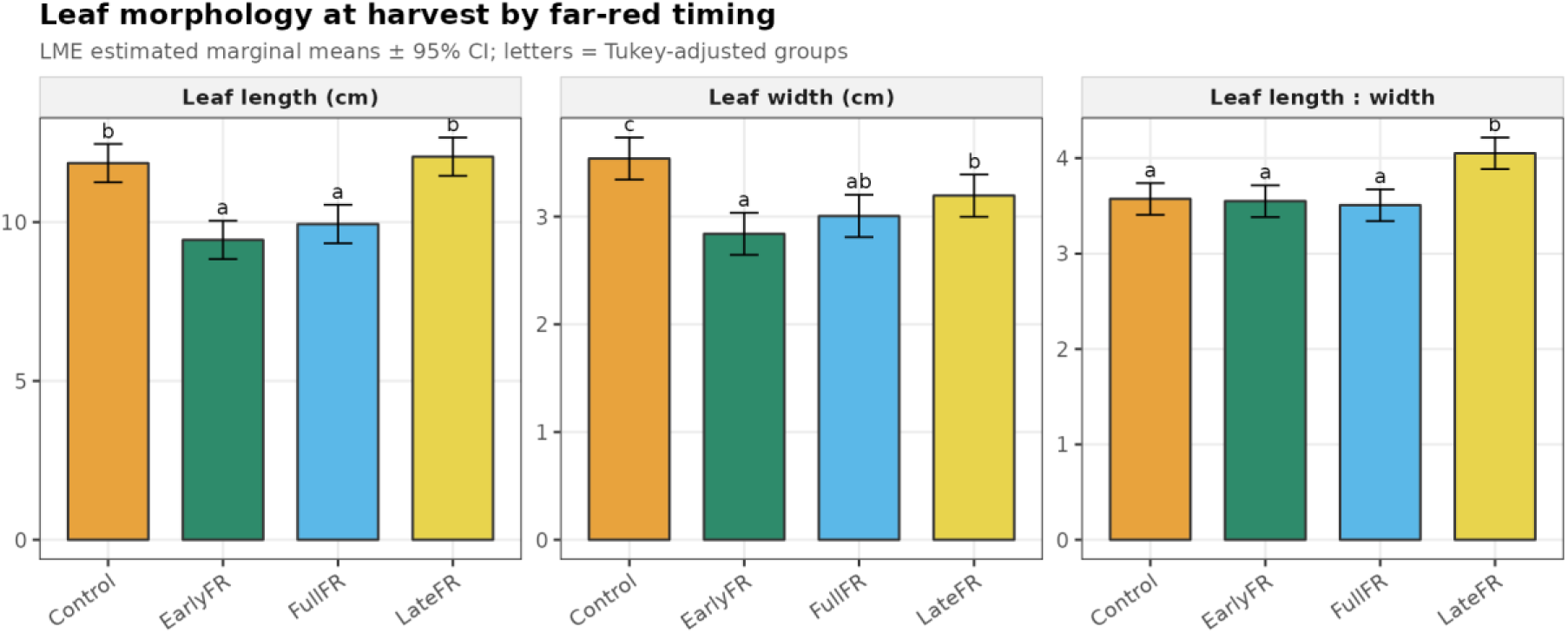
Leaf morphology at harvest (T2): (a) leaf length, (b) leaf width and (c) leaf length-to-width ratio by far-red timing. Values are LME estimated marginal means ± 95 % CI; letters indicate Tukey-adjusted groups. Early and continuous far-red reduced leaf size, whereas late far-red produced the highest length-to-width ratio.

### Chlorophyll

Total chlorophyll differed among treatments (F(3,72) = 17.3, p < 0.001), with no significant cultivar effect (p = 0.68) or treatment × cultivar interaction (p = 0.35). EarlyFR produced the highest total chlorophyll and was significantly greater than the other treatments when treatment effects were isolated (Fig. 7a). Chlorophyll a:b ratio also differed by treatment (F(3,72) = 16.8, p < 0.001) but not by cultivar. Control had the highest a:b ratio and EarlyFR the lowest, with FullFR and LateFR intermediate (Fig. 7b). The reduced a:b ratio under EarlyFR indicates relative enrichment of chlorophyll b, consistent with a shade response. These findings only partly support the hypothesis of neutral or negative FR effects on chlorophyll, because EarlyFR increased total chlorophyll.

**Figure 7.**
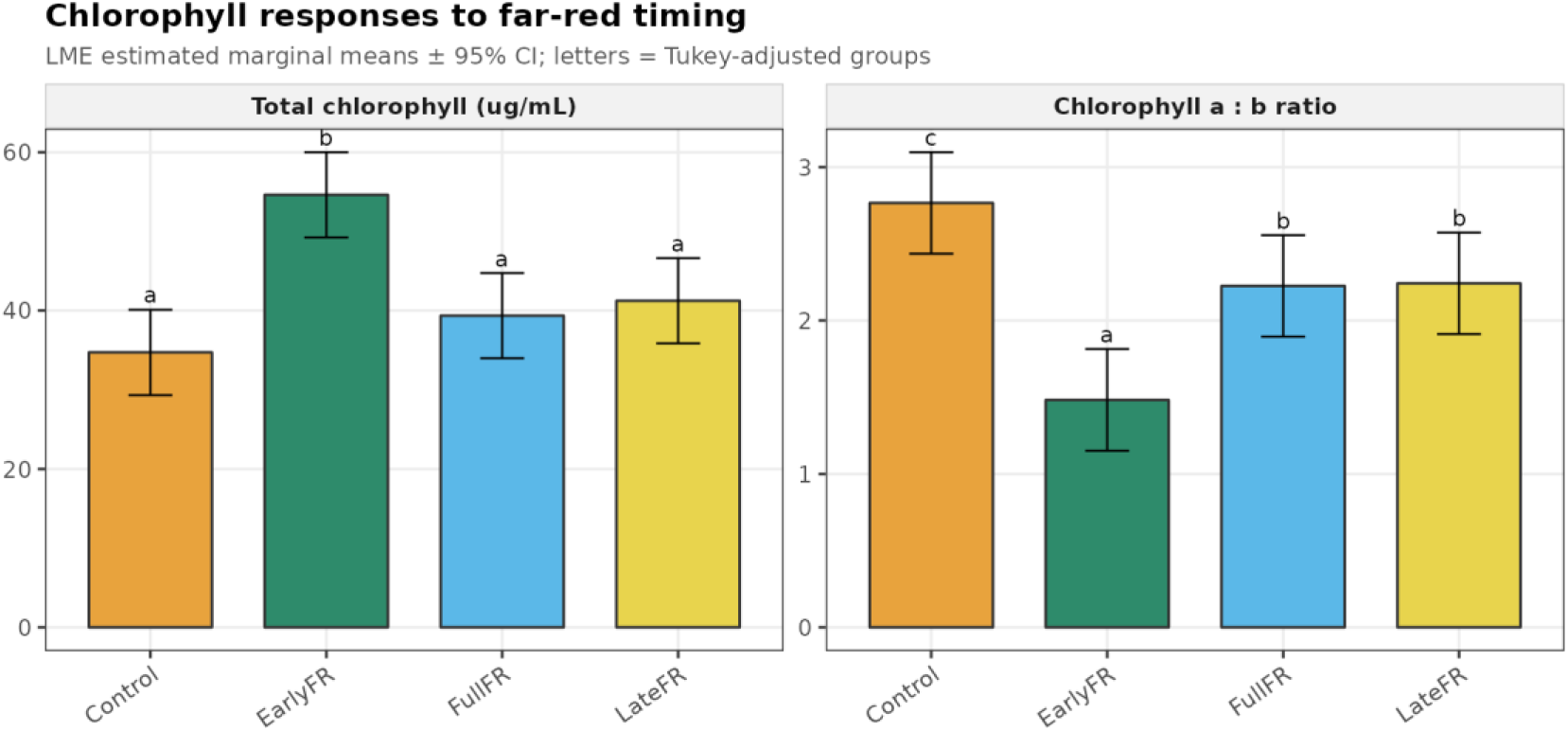
(a) Total chlorophyll and (b) chlorophyll a:b ratio by treatment. Values are LME estimated marginal means ± 95 % CI; letters indicate Tukey groups. Early far-red increased total chlorophyll and reduced chlorophyll a:b ratio, consistent with chlorophyll-b enrichment under an early shade signal.

### Suitability ranking for vertical farming

Treatment rankings based on high marketable yield and low flowering placed LateFR first and EarlyFR last, with LateFR outperforming Control and differing significantly from EarlyFR and FullFR (Fig. 8a). A composite cultivar ranking based on high yield, low flowering and high chlorophyll identified Santa Cruz as the strongest overall cultivar. Although Santa Cruz ranked fourth for raw fresh weight, it combined low flowering incidence, high chlorophyll and compact growth (Fig. 8b). High-yielding but bolt-prone cultivars, including Amador, Harp and Rubino, ranked lower once flowering liability was included.

**Figure 8.**
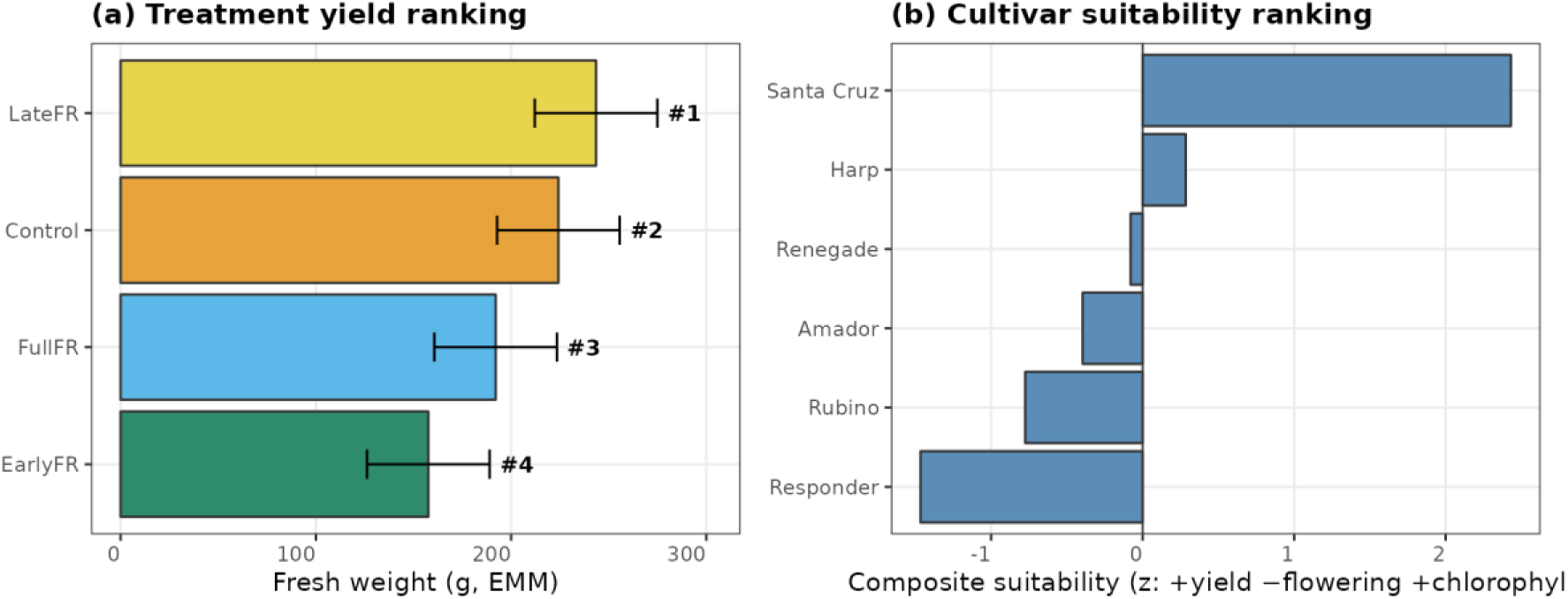
Suitability for vertical-farm production. (a) Treatment ranking by marketable fresh weight (estimated marginal means; #1 = best). (b) Composite cultivar suitability based on standardised positive yield, negative flowering and positive chlorophyll scores; Santa Cruz ranked highest overall.

### Summary of statistical models

**Table 3** provides a consolidated summary of fixed-effect tests. Treatment was significant for every measured response. Cultivar was significant for all traits except chlorophyll. The treatment × cultivar interaction was significant for dry weight, leaf morphology, height and flowering, but not for fresh weight or chlorophyll.

To indicate how strongly each factor shaped the measured responses, the explained variation was partitioned among treatment, cultivar and their interaction (Table 4). Cultivar accounted for the largest share of the factorial variation in most traits, including dry weight (64 %), the leaf dimensions (leaf width 82 %, leaf length:width ratio 89 %), plant height (92 %) and flowering incidence (79 %), whereas far-red timing dominated only the two chlorophyll traits (71–77 %). Marketable fresh weight was the most evenly partitioned trait (cultivar 48 %, treatment 30 %, interaction 22 %). The treatment × cultivar interaction was a minority component in every trait, ranging from 2 % for height to 23 % for dry weight, yet it remained statistically significant for dry weight, leaf morphology, height and flowering (Table 3). The interaction therefore carries agronomic weight for cultivar-specific light management that is not reflected in its share of variance alone, and larger trials would allow this genotype × environment × management component to be estimated with greater precision.

### Broad-sense heritability and projected genetic gain

Because cultivar was represented by a fixed panel of six commercial varieties, we summarised the repeatability of each trait as a broad-sense heritability on an entry-mean basis, estimated from the cultivar F-ratios of the split-plot models as H² = 1 − 1/F (Falconer and Mackay, 1996; Holland et al., 2003; Table 5). Heritability was high for the agronomic and architectural traits — marketable fresh weight (H² = 0.88), dry weight (0.93), the leaf dimensions (0.94–0.99) and plant height (0.99) — but was not estimable for the two chlorophyll traits, for which cultivar was non-significant. These values describe the among-cultivar signal expressed across the four far-red regimes and are therefore an upper bound on the heritability that would be realised when selecting on single plots; we use them to anchor, rather than to fix, the projections below. Using the breeder’s equation in its response-to-selection form (ΔG = i r σ_G), projected gain increased with heritability and selection intensity as expected (Supplementary Fig S1a). Notably, at the heritabilities estimated here, simple phenotypic selection on marketable yield matched or exceeded the gain from a moderate-accuracy genomic-prediction scenario, and genomic prediction became the more efficient route only at lower heritabilities (Supplementary Fig S1a). When the same expression was placed in its extended form (ΔG = i r_MET r_MET,TPE σ_G / L; Cooper et al., 2023), the compressed breeding generation interval achievable indoors (of the order of weeks to a few months, against a single field cycle per year) and the close correspondence between the trial and the target production environment of a vertical farm (r_MET,TPE → 1) together projected substantially higher annual gain than a comparable open-field multi-environment trial (Supplementary Fig S1b).

## Discussion

### Far-red timing as a management lever

This study demonstrates that the developmental window in which supplemental FR is applied, rather than its cumulative dose, governs the agronomic outcome in spinach; indeed LateFR delivered more total far-red than EarlyFR (21.3 vs 13.5 mol m⁻²) yet patterned with the unsupplemented Control. LateFR increased marketable fresh weight, increased leaf length-to-width ratio and reduced bolting, whereas EarlyFR and FullFR reduced yield and increased flowering. These findings shift the interpretation of FR from a fixed spectral component to a stage-specific management factor. They may also help reconcile our results with Skabelund, Langenfeld and Bugbee (2025), who found no morphological response to FR fraction in spinach. If FR sensitivity is developmentally gated, experiments that vary FR fraction while holding the exposure window constant may fail to detect responses that emerge when timing is varied directly.

### Flowering and bolting

Early FR increased flowering, consistent with the role of low R:FR as a signal of neighbour proximity during early development. Under such conditions, shade-avoidance pathways can promote stem elongation and earlier reproductive transition before canopy closure (Franklin & Whitelam, 2005; Casal, 2012; Tan et al., 2022). Continuous FR maintained this signal and also increased flowering. In contrast, plants exposed to FR only later in development appeared to respond primarily through architectural adjustment, including greater height and altered leaf proportions, rather than through a comparable increase in reproductive commitment. The low blue-to-red background may have further favoured PIF-mediated elongation and flowering under FR treatments (Casal, 2012; Park & Runkle, 2019). These results are consistent with the wider literature on FR and flowering, but add a temporal component that has been little explored in spinach. They should also be interpreted in relation to photoperiod and temperature, both of which strongly influence spinach flowering: the 12 h photoperiod and 24 °C temperature used here are close to thresholds associated with reproductive transition in susceptible genotypes (Chun et al., 2000; Isaza et al., 2025), and phytochrome-B responses can be temperature-sensitive (Halliday et al., 2003).

### Biomass and resource allocation

Fresh weight was inversely related to flowering response. Bolting redirects growth toward stem and reproductive structures and away from harvestable leaves, reducing both yield and potential photosynthetic return (de Wit et al., 2018; Amini et al., 2025). The largest timing effects occurred in bolt-resistant cultivars because these genotypes flowered little under Control and LateFR but were induced to bolt by EarlyFR. In contrast, bolt-prone cultivars flowered under most conditions and were therefore less responsive to changes in timing. Because biomass was recorded as total aerial mass, this study could not partition leaf, stem and flower biomass directly. Future experiments should include organ-level biomass partitioning to quantify allocation shifts. Nevertheless, the practical result is clear: FR applied after early canopy establishment can increase harvestable yield, and its contribution to photosynthetic efficiency may reduce lighting requirements when timed appropriately (Zhen & Bugbee, 2020; Farhangi et al., 2025).

### Leaf morphology and physiology

The increase in leaf length-to-width ratio under FR, especially under LateFR, is consistent with a shade-avoidance elongation response mediated by phytochrome-B and downstream SAR regulators (Galvão et al., 2019; Paik & Huq, 2019). Because leaf and petiole were measured together, the present data cannot distinguish lamina expansion from petiole elongation and therefore cannot directly test specific-leaf-area effects reported in other spinach studies (Lőrinc et al., 2019; Skabelund et al., 2025). The chlorophyll response was less typical. EarlyFR increased total chlorophyll and reduced chlorophyll a:b ratio, whereas FR often reduces chlorophyll content in other species. One possible explanation is that early FR promoted development of chlorophyll-b-rich light-harvesting antennae while maintaining chlorophyll a, leading to a higher total chlorophyll signal (Voitsekhovskaja & Tyutereva, 2015; Chen et al., 2021). Continuous FR may have attenuated or desensitised this response. Because chlorophyll was measured on standard leaf samples rather than per unit mass, these results should be interpreted as relative differences. Comparable far-red-driven elongation responses have recently been shown to differ markedly among genotypes in other crops; in quinoa, for example, supplemental far-red increased stem elongation in all genotypes tested but to differing degrees (Gordillo-Romero et al., 2026).

### Synthesis: a G×E×M recommendation

Together, the results define a G×E×M interaction in which the effect of FR enrichment depends on both application timing and cultivar bolting tolerance. For vertical-farm production, two management implications follow. First, supplemental FR should be applied after the early growth stage rather than throughout production or during early establishment. Second, cultivar choice should be made in relation to FR timing. Santa Cruz ranked highest overall because it combined low flowering, high chlorophyll and compact growth, whereas high-fresh-weight but bolt-prone cultivars ranked lower when flowering was penalised. The same principle may be relevant beyond vertical farming, because cultivar selection and managed light environments could help extend harvest windows under increasingly variable production conditions. Genotype-dependent responses to light quality of this kind are increasingly reported across crops, including a recent demonstration in quinoa that supplemental far-red, red and blue light altered growth and stress responses in a strongly genotype-specific manner (Gordillo-Romero et al., 2026), reinforcing the value of evaluating spectral management within a genotype-aware framework.

### From management response to genetic gain

The genotype × management interaction documented above (Section 3.3; Table 4) has a direct quantitative-genetic reading. Within the breeder’s equation, the response to selection for marketable yield depends on the genetic variance and heritability that are actually expressed, and both are conditioned by the management regime. Early far-red compressed the cultivar panel toward bolting and yield loss, whereas late far-red raised the mean and separated cultivars according to their intrinsic bolting tolerance; management therefore determines which genetic differences are visible to selection. Optimising genotype and management jointly — rather than fixing a light recipe and then choosing a cultivar, or the reverse — is thus expected to increase the achievable rate of gain, which is the central argument for embedding G×E×M within predictive-breeding frameworks (Cooper et al., 2023; Powell et al., 2026). Controlled-environment vertical farms are unusually well suited to this approach. Because the trial environment is the production environment, the alignment between the evaluation environment and the target population of environments approaches unity, removing much of the genotype-by-environment uncertainty that constrains gain in field programmes (Powell et al., 2026); and because growth is rapid and season-independent, several generations can be advanced per year rather than the single annual cycle typical of field production. Treating far-red timing as a heritability-modifying management variable, and selecting cultivars within the chosen regime, therefore offers a tractable route to accelerate the improvement of indoor leafy crops. We did not estimate narrow-sense heritability or genetic gain directly, because the six cultivars were a fixed commercial panel rather than a segregating population; the projections in Fig. 8 are intended as a quantitative framing of this opportunity and a guide to the larger, pedigreed trials that would be required to realise it.

### Limitations

Several limitations qualify these conclusions. First, height at T2 partly reflects bolting and should not be interpreted as an independent vegetative-growth trait. Second, total aerial biomass was measured without separating leaf, stem and reproductive tissues, limiting inference about allocation. Third, chlorophyll values were relative rather than mass-based and were derived from two leaves per tray. Fourth, FR was supplied additively; therefore, total photon flux differed between FR and non-FR phases even though PAR was constant. The observed effects cannot be attributed exclusively to FR signalling independent of a small increase in total photons. We addressed this dose confound directly: because LateFR carried a larger far-red dose than EarlyFR yet produced the opposite response, the outcomes cannot be explained by cumulative photon dose, and an estimated phytochrome-state metric outperformed a dose model (Section 3.4). The phytochrome photostationary state was nonetheless calculated from programmed light recipes rather than measured spectra (channel cross-sections calibrated to reproduce published Sager-based values; Lanoue et al., 2022), the early competence window was specified a priori and is partly confounded with the EarlyFR treatment, and the early-window deficit took only two distinct values, so the cultivar reaction norms represent two-level contrasts rather than continuous dose–response relationships. Finally, replication was modest (five trays per cultivar per treatment), consistent with the non-significant fresh-weight interaction. These limitations do not alter the main timing result but identify priorities for follow-up experiments.

## Conclusion

Far-red timing altered the developmental and agronomic response of spinach grown in a controlled environment vertical-farm system. Late FR increased marketable fresh weight, increased leaf length-to-width ratio and reduced bolting, whereas early FR increased flowering and reduced yield. Early FR also increased total chlorophyll and reduced chlorophyll a:b ratio, indicating that physiological as well as morphological responses depended on timing. The magnitude of these effects varied among cultivars and was greatest in cultivars with lower baseline bolting. We conclude that supplemental FR should be applied after early growth and that cultivar selection and FR timing should be optimised jointly as a G×E×M decision. Framing these timing effects within the breeder’s equation further suggests that vertical farms, in which the trial and production environments coincide and breeding generation intervals can be short, are a favourable setting in which to optimise genotype and far-red management jointly for genetic gain (Powell et al., 2026). Future work should partition biomass by organ, quantify chlorophyll per unit mass, equalise total photon flux and test FR timing under contrasting photoperiod and temperature regimes.

## Author Contributions

Christopher McGovern, Hadi Aliki, and Rajiv Sharma conceived and designed the study. Christopher McGovern and Milena Adrio conducted the experiments and collected the data. Rajiv Sharma and Wayne Powell led the data analysis and interpretation, with contributions from Christopher McGovern, Milena Adrio and Ruth Vichos. Rajiv Sharma supervised the research, with support from Hadi Aliki and Ruth Vichos. Rajiv Sharma led the writing of the original manuscript, with contributions from Christopher McGovern. Wayne Powell, Christopher McGovern, Milena Adrio, Hadi Aliki, and Ruth Vichos reviewed and edited the manuscript. All authors read and approved the final manuscript.

## Acknowledgements

The authors gratefully acknowledge the support of Scotland’s Rural College (SRUC) through core funding for the vertical farming and greenhouse facilities. We also wish to thank the technical staff for their invaluable assistance with vertical-farm operations and data collection.

## Data Availability Statement

All data generated or analyzed during this study are included in this published article and its supplementary information files.

## Supplementary Table Legends

**Supplementary Table S1. Phytochrome photostationary state (PSS) predictors.** Experimental treatment details including Far-red (FR) window timing, cumulative FR dose (mol m⁻²), and associated PSS values (*φ_FR_*__*on*_, *φ_FR_*__*off*_) and Early-window Pfr-deficit (*φ_d_*) metrics.

**Supplementary Table S2. PSS calibration anchors.** Validation data comparing target PSS values derived from Sager et al. benchmarks against fitted PSS values across various source treatments.

**Supplementary Table S3. Spectral cross-sections and channel-specific PSS.** Summary of spectral sensitivity parameters (*σ_r_*, *σ_fr_*) and calculated PSS for the blue, green, red, and far-red light channels.

**Supplementary Table S4. Cultivar bolting resistance and flowering incidence.** Bolting resistance scores and flowering incidence (%) for six spinach cultivars (Amador, Harp, Renegade, Responder, Rubino, Santa Cruz) across four light treatments: Control, LateFR, EarlyFR, and FullFR.

**Supplementary Table S5. Statistical model comparison for bolting response.** Comparison of AIC values for models predicting bolting incidence, highlighting the model performance difference (ΔAIC = 441) between the Early-window Pfr-deficit and Far-red dose models.

